# Closed-loop control of sensorimotor aperiodic neural dynamics can manipulate excitatory-inhibitory balance in humans

**DOI:** 10.64898/2026.11.09.765129

**Authors:** Satoshi Mutsuda, Seitaro Iwama, Junichi Ushiba

## Abstract

A ubiquitous property of physical and biological systems is scale-free dynamics in which spectral power decreases as a function of frequency, showing a 1/f spectral slope. The 1/f slope of neural power spectra has been proposed as a marker of cortical excitation-inhibition (E/I) balance, based on pharmacological experiments and computational models of synaptic activity in cortical networks. However, whether or not the spectral slope in the human population recordings, influenced by biophysical or experimental factors, can be utilized as an index of cortical E/I is debatable. Here, we show that this slope can be volitionally modulated and that its modulation is accompanied by changes in motor cortical activity reflecting E/I balance. To this end, we used a closed-loop paradigm in which real-time changes in the sensorimotor 1/f slope during neurofeedback trigger transcranial magnetic stimulation to probe intracortical circuits in the primary motor cortex. We found lower facilitatory effects in the steeper 1/f state than in both the flatter state and at rest, despite no significant between-state differences in 1/f-adjusted band power. These findings establish the physiological relevance of 1/f modulation with cortical states and support the contribution of intracortical E/I-related processes to the aperiodic dynamics.

## Introduction

A ubiquitous property of physical and biological systems is scale-free dynamics in which spectral power decreases as a function of frequency, referred to as a 1/f-like relationship^1,2^. This characteristic spectral decay has been interpreted as a statistical signature of collective dynamics and self-organized processes^3^. In recent years, similar broadband, scale-free components have been identified in human electrophysiological recordings. In addition to the periodic population activity, the slope of this aperiodic component, parameterized as the aperiodic exponent (χ) in the relationship *P*(*f*) ∝ 1⁄*f***^z^**^4^, varies systematically across behavioral and physiological states, suggesting that it may index large-scale cortical population dynamics^5,6^.

The spectral slope in neural power spectra is thought to partly reflect the balance between excitatory and inhibitory (E/I) synaptic activity as well as network time constants^7–9^. Previous studies have typically measured electroencephalogram (EEG) during task performance or following pharmacological interventions and examined correlations between spectral features and behavioral or physiological indices^6,10^. While these findings suggest that the 1/f slope may relate to E/I balance or behavioral state, it remains unclear whether the task-related modulation of aperiodic activity is driven by changes in E/I balance. Since there are possible alternative modulators of aperiodic activity^11–13^, this calls for direct experimental validation that manipulates intrinsic aperiodic activity to probe the instantaneous excitatory and inhibitory circuit profile.

Here, we sought to determine whether the aperiodic exponent of human EEG reflects motor cortical circuit dynamics. To this end, we developed a closed-loop EEG-triggered transcranial magnetic stimulation (TMS) paradigm that combined real-time neurofeedback of the sensorimotor aperiodic exponent with physiological probing of the primary motor cortex (M1). TMS applied to M1 offers a unique opportunity to address this question. M1 possesses well-characterized recurrent circuitry and direct corticospinal output pathways^14–16^ and its intracortical inhibitory and facilitatory processes can be quantified using paired-pulse protocols^17,18^. Therefore, experimentally modulating the aperiodic exponent in M1 and probing corresponding changes in motor cortical physiology provides a means to test whether the spectral slope reflects measurable circuit-level dynamics.

Participants were guided to volitionally induce flatter or steeper spectral slopes relative to rest, and TMS was triggered over M1 contingent on these real-time changes (Fig. 1a). This closed-loop paradigm allowed us to quantify corticospinal excitability using single-pulse motor-evoked potentials (MEPs), together with intracortical inhibition and facilitation using paired-pulse TMS. If the aperiodic exponent indexes underlying synaptic population dynamics, voluntarily induced shifts in spectral slope would change TMS-derived measures of motor cortical circuit function.

**Figure 1.**
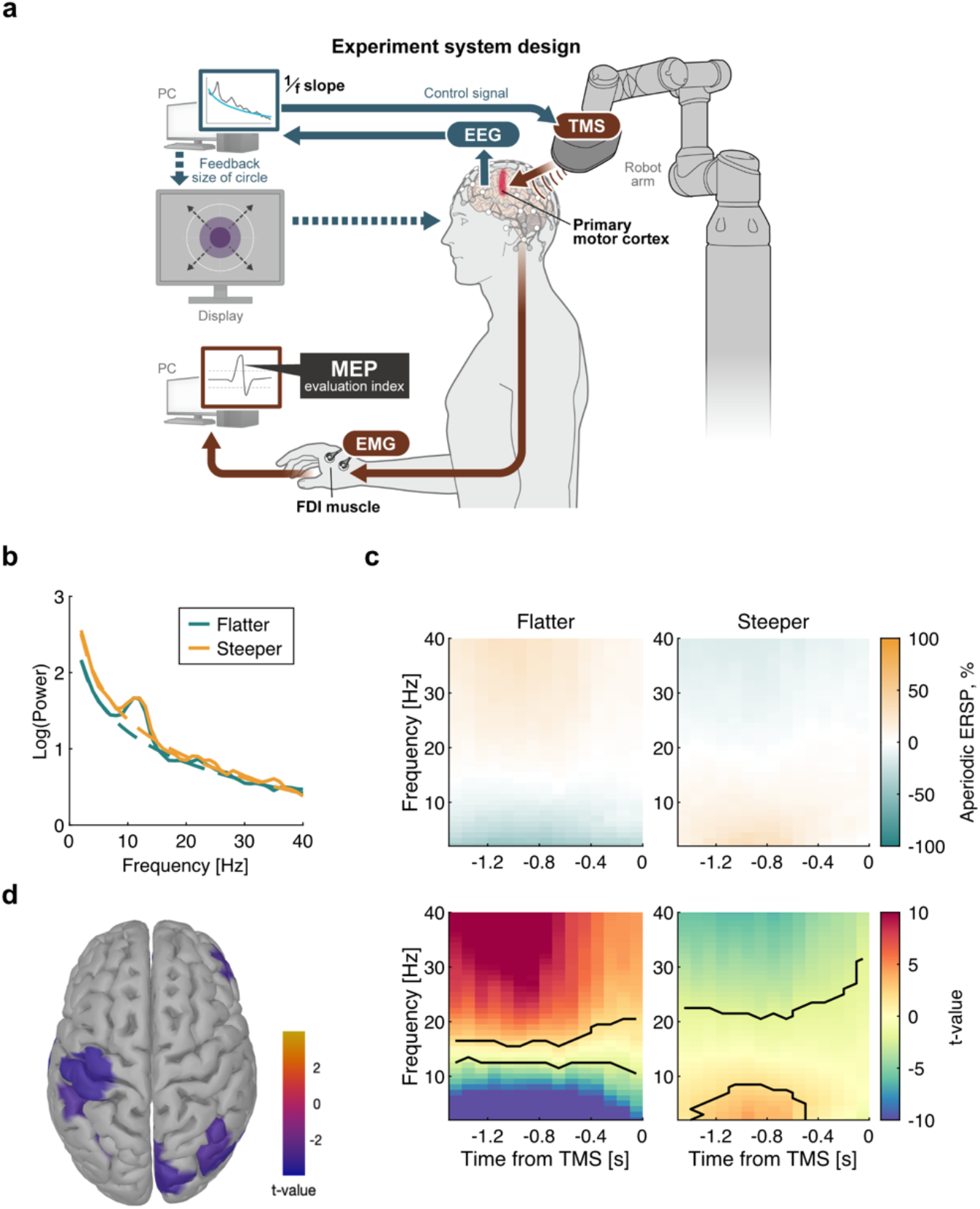
Bidirectional modulation of aperiodic activity across sensor and source space. (a) Schematic of the closed-loop EEG–TMS experimental system. The aperiodic exponent was estimated online from sensorimotor EEG activity and presented as visual neurofeedback. TMS was delivered over the primary motor cortex (M1) contingent on modulation of the aperiodic exponent, and motor-evoked potentials (MEPs) were recorded from the first dorsal interosseous (FDI) muscle using electromyography (EMG). (b) Representative power spectral density (PSD) profiles during the flatter and steeper neurofeedback conditions, illustrating the condition-dependent difference in broadband spectral slope. (c) Time–frequency representations of aperiodic power changes preceding TMS. Upper panels show aperiodic ERSP relative to the resting-state condition for the flatter and steeper conditions. Lower panels show t-value maps from cluster-based permutation tests against rest. Black contours indicate clusters reaching cluster-level significance (*p* < 0.05). (d) Cortical distribution of significant condition-dependent differences in the aperiodic exponent. Colors indicate t values for the flatter-minus-steeper contrast in parcels surviving Benjamini–Hochberg FDR correction across 100 cortical parcels; nonsignificant parcels are shown without color coding.

## Results

Participants underwent simultaneous EEG–TMS recordings under three experimental conditions. In the two neurofeedback conditions, they were guided to self-regulate the aperiodic exponent toward either a flatter or steeper state, with TMS delivered whenever modulation reached 15% according to one of three stimulation protocols. In the resting-state condition, visual feedback was presented without neurofeedback, and TMS was delivered repeatedly. All procedures were well tolerated, with no adverse events reported.

### Bidirectional modulation of aperiodic activity across sensor and source space

We first sought to determine whether neurofeedback could induce task-related modulation of aperiodic activity in the targeted sensorimotor cortex. To characterize the spectral and spatial expression of this modulation, we examined power spectral density (PSD) profiles, aperiodic time–frequency dynamics, and their cortical source distribution. Representative PSD profiles illustrated the spectral dissociation between the two neurofeedback states (Fig. 1b). The broadband spectral slope differed between conditions, whereas the alpha-band peak appeared comparable.

To visualize which frequency ranges carried the power changes associated with modulation of the aperiodic exponent, we analyzed time–frequency maps reconstructed from the aperiodic component alone (Fig. 1c). Event-related spectral perturbation of aperiodic activity (Aperiodic ERSP) showed opposing frequency-dependent patterns across the two neurofeedback conditions. Relative to rest, the flatter condition showed significant negative and positive clusters at lower (2–13 Hz) and higher frequencies (16– 40 Hz), respectively, whereas the steeper condition showed the opposite pattern, with a significant positive cluster at lower frequencies (2–8 Hz) and a negative cluster at higher frequencies (21–40 Hz) (Cluster-based permutation tests, *p*_cluster_ < 0.05; Fig. 1c). Thus, the two neurofeedback-induced cortical states were characterized by opposing broadband power changes across lower and higher frequencies.

Having established bidirectional aperiodic modulation in the sensor space, we next examined whether this condition-dependent effect was also evident in source space (Fig. 1d). Source reconstruction revealed a spatially widespread condition-dependent pattern, with the flatter-minus-steeper contrast showing the same direction in 99 of 100 cortical parcels. Fifteen parcels showed significant differences after false discovery rate (FDR) correction, including the left somatomotor parcel associated with the targeted C3 electrode (*t*(28) = −3.29, *p* = 0.003, *p_FDR_* = 0.034, *d_z_* = −0.61). Together, these findings indicate that the condition-dependent modulation observed at the sensor level was also evident in source space, including the targeted sensorimotor region.

### Bidirectional modulation of the aperiodic exponent

The opposing broadband spectral patterns observed between the two neurofeedback states suggest that they may arise from modulation of the aperiodic exponent. We therefore tested whether the experimentally induced spectral changes were quantitatively captured by changes in the aperiodic exponent. To quantify the modulation depth of the cortical spectral slope, we calculated Δχ as the relative change in the aperiodic exponent from the condition-specific resting baseline (Fig. 2a). We then compared Δχ between trials in which the online Δχ reached the target threshold (success) and those in which it did not (failure) for each target direction. A one-way repeated-measures ANOVA revealed a significant main effect of state on Δχ (*F*(1.79, 50.06) = 126.9, *p* < 0.001, ηp² = 0.82; Fig. 2a). Post hoc comparisons showed that success-flatter trials had significantly lower Δχ than failure-flatter trials (*t*(28) = 11.50, corrected-*p* < 0.001, *d* = 2.82), whereas success-steeper trials had significantly higher Δχ than failure-steeper trials (*t*(28) = 13.56, corrected-*p* < 0.001, *d* = 2.83). Success-steeper trials were also significantly higher than success-flatter trials (*t*(28) = 20.82, corrected-*p* < 0.001, *d* = 3.53), confirming a robust target-dependent separation in the offline aperiodic exponent measure. Thus, successful neurofeedback was associated with bidirectional modulation of Δχ in the instructed direction, validating the online classification of flatter and steeper feedback success.

**Figure 2.**
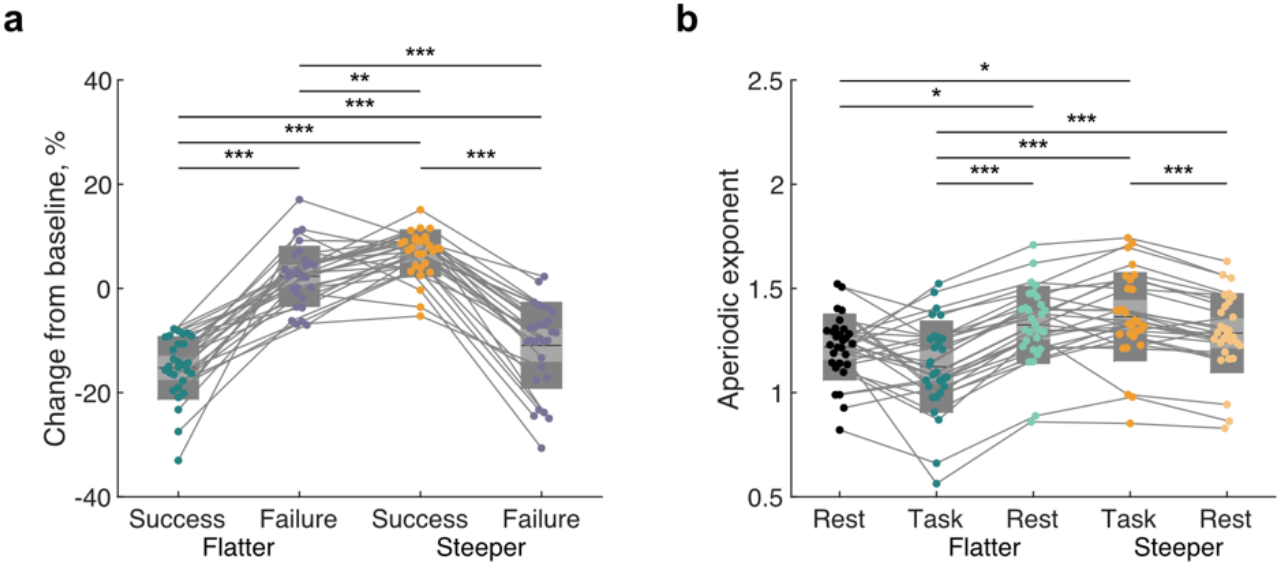
Bidirectional modulation of the pre-TMS aperiodic exponent. (a) Trial-wise changes in the aperiodic exponent (Δχ) across success and failure trials in the flatter and steeper conditions. Δχ was calculated as (χ_task_ – χ_rest_) / χ_rest_, where χ_task_ and χ_rest_ represent the aperiodic exponent during the task and rest periods within each trial, respectively. (b) Raw aperiodic exponent during success trials in the flatter and steeper conditions, their corresponding condition-specific baseline periods, and an independent eyes-open resting-state recording. In all panels, dots represent individual participants, gray lines connect measurements from the same participant, and box plots indicate the mean ± 1 SD (dark gray) and the 95% confidence interval of the mean (light gray). Black horizontal lines indicate the group means. Statistical significance is indicated as \**p* < 0.05, \*\**p* < 0.01, and \*\*\**p* < 0.001; n.s., not significant.

To determine whether the observed bidirectional effect depended on baseline normalization, we next examined the raw aperiodic exponent (Fig. 2b). Successful flatter and steeper trials were evaluated against both the condition-specific baselines used for Δχ calculation and an independent 1-min eyes-open resting-state recording. A one-way repeated-measures ANOVA revealed a significant main effect of state on the raw aperiodic exponent (*F*(1.77, 49.45) = 24.25, *p* < 0.001, ηp² = 0.46). Post hoc comparisons showed that successful flatter trials were significantly lower than the corresponding flatter baseline (*t*(28) = 17.38, corrected-*p* < 0.001, *d* = 1.04), whereas successful steeper trials were significantly higher than the corresponding steeper baseline (*t*(28) = 7.24, corrected-*p* < 0.001, *d* = 0.41). Successful steeper trials were also significantly higher than successful flatter trials (*t*(28) = 11.30, corrected-*p* < 0.001, *d* = 1.24), indicating bidirectional modulation of the raw aperiodic exponent.

Relative to the independent eyes-open resting state, the condition-specific baseline for the flatter condition showed a significantly higher exponent (*t*(28) = 3.20, corrected-*p* = 0.035, *d* = 0.55), whereas the baseline for the steeper condition did not differ significantly from rest (*t*(28) = 1.87, corrected-*p* = 0.727, *d* = 0.34). Successful steeper trials showed significantly higher exponent values than the independent resting state (*t*(28) = 3.67, corrected-*p* = 0.010, *d* = 0.75), whereas successful flatter trials did not differ significantly from rest (*t*(28) = 2.35, corrected-*p* = 0.263, *d* = 0.49). Thus, although the aperiodic exponent shifted in the instructed direction relative to the condition-specific baseline in both neurofeedback conditions, only the steeper condition showed a significant deviation from the independent resting state during successful trials.

### Dissociating aperiodic and periodic contributions to conventional band-power modulation

We next investigated whether this modulation occurred independently of changes in periodic oscillatory activity. Because band-power measures derived from the original power spectra contain both periodic and aperiodic contributions, we first quantified conventional ERSP in theta, alpha, and beta bands using the original spectra and then repeated the analysis after removing the fitted aperiodic component to isolate periodic activity. Conventional ERSP differed significantly between the flatter and steeper states in the theta band (*t*(28) = 2.85, *p* = 0.008, *d*_z_ = 0.53) and beta band (*t*(28) = 3.38, *p* = 0.002, *d*_z_ = 0.63), whereas no significant difference was observed in the alpha band (*t*(28) = 0.32, *p* = 0.748, *d*_z_ = 0.06; Fig. 3a). These opposing low-and high-frequency differences were consistent with the spectral pattern expected from the experimentally induced modulation of the aperiodic slope.

**Figure 3.**
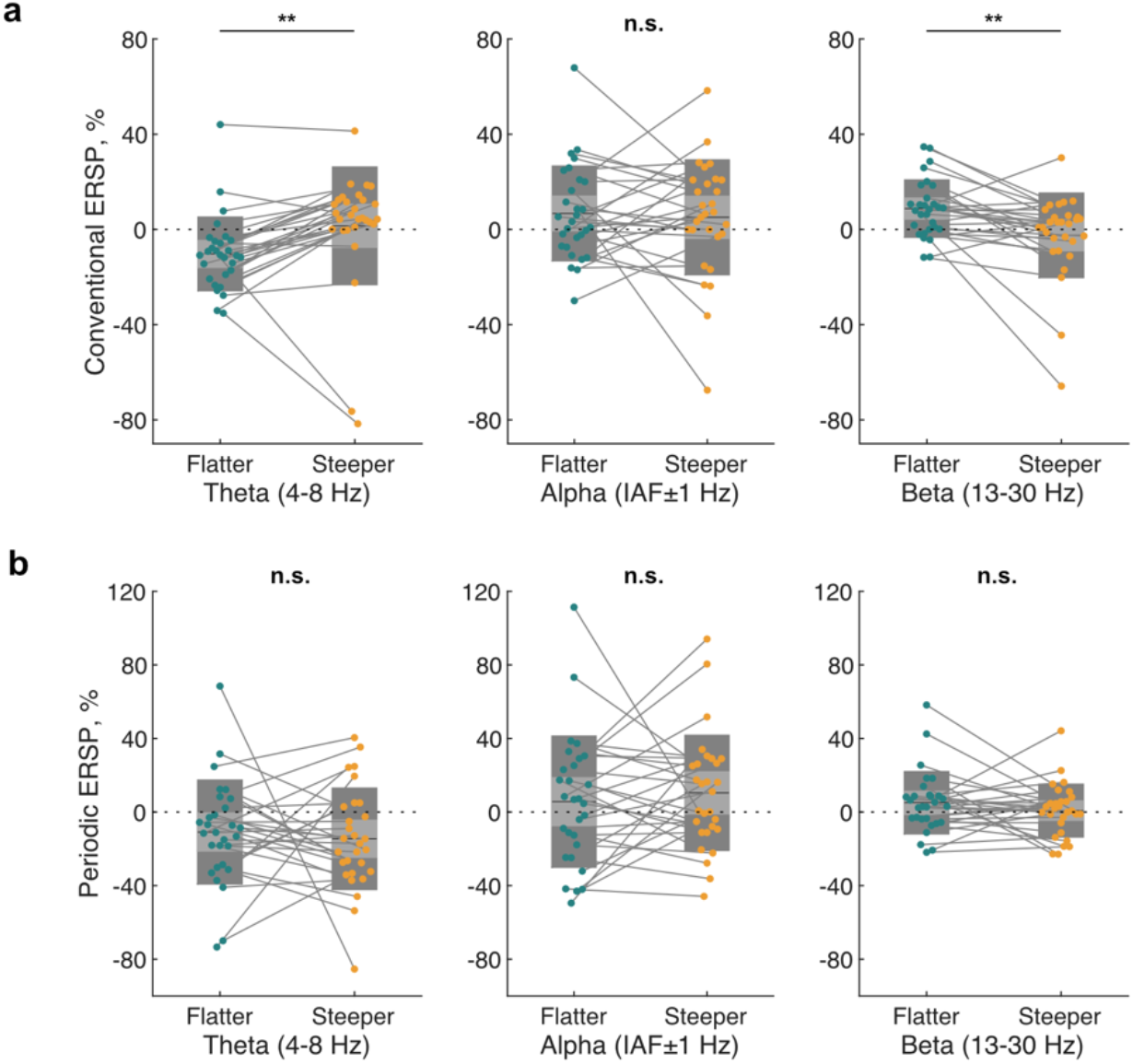
Aperiodic exponent modulation is associated with conventional but not periodic band-power differences. (a) Conventional ERSP in the theta (4–8 Hz), individualized alpha (IAF ± 1 Hz), and beta (13–30 Hz) bands during the Flatter and Steeper neurofeedback states. Significant condition differences were observed in the theta and beta bands, but not in the alpha band. (b) Periodic ERSP within the same frequency bands after removal of the fitted aperiodic component. No significant condition differences were observed in periodic theta-, alpha-, or beta-band activity. Dots represent individual participants, with paired observations connected by gray lines. Gray boxes indicate the mean ± 1 SD (dark gray) and the 95% confidence interval of the mean (light gray). Black horizontal lines indicate the group means. The horizontal dotted line indicates no change relative to baseline. Statistical comparisons were performed using paired-sample *t*-tests. \*\**p* < 0.01; n.s., not significant.

To isolate changes in periodic activity, we applied the identical analysis to the flattened spectra obtained by subtracting the fitted aperiodic component from the original PSD (Fig. 3b). After accounting for the aperiodic background, none of the three frequency bands showed a significant difference between the flatter and steeper states (theta: *t*(28) = 0.47, *p* = 0.646, *d*_z_ = 0.09; alpha: *t*(28) = 0.66, *p* = 0.518, *d*_z_ = 0.12; beta: *t*(28) = 1.17, *p* = 0.254, *d*_z_ = 0.22). Notably, the significant theta-band difference observed in conventional ERSP was no longer present after aperiodic correction, and the numerical direction of the condition difference was reversed (flatter: −10.88 ± 5.24%; steeper: −14.53 ± 5.12%), although this difference was not significant. Thus, the bidirectional modulation of the aperiodic exponent was achieved without a significant condition-dependent modulation of periodic activity within the theta, alpha, or beta bands.

### Direction-specific differences in intracortical facilitation across neurofeedback-induced aperiodic states

Although we have confirmed that neurofeedback induced distinct aperiodic states in the absence of measurable differences in periodic activity, this raised the question of whether these states were also accompanied by distinct facilitatory and inhibitory circuit profiles in the motor cortex. To address this question, we compared corticospinal excitability, intracortical inhibition, and intracortical facilitation across the flatter, steeper, and rest states (Fig. 4a).

**Figure 4.**
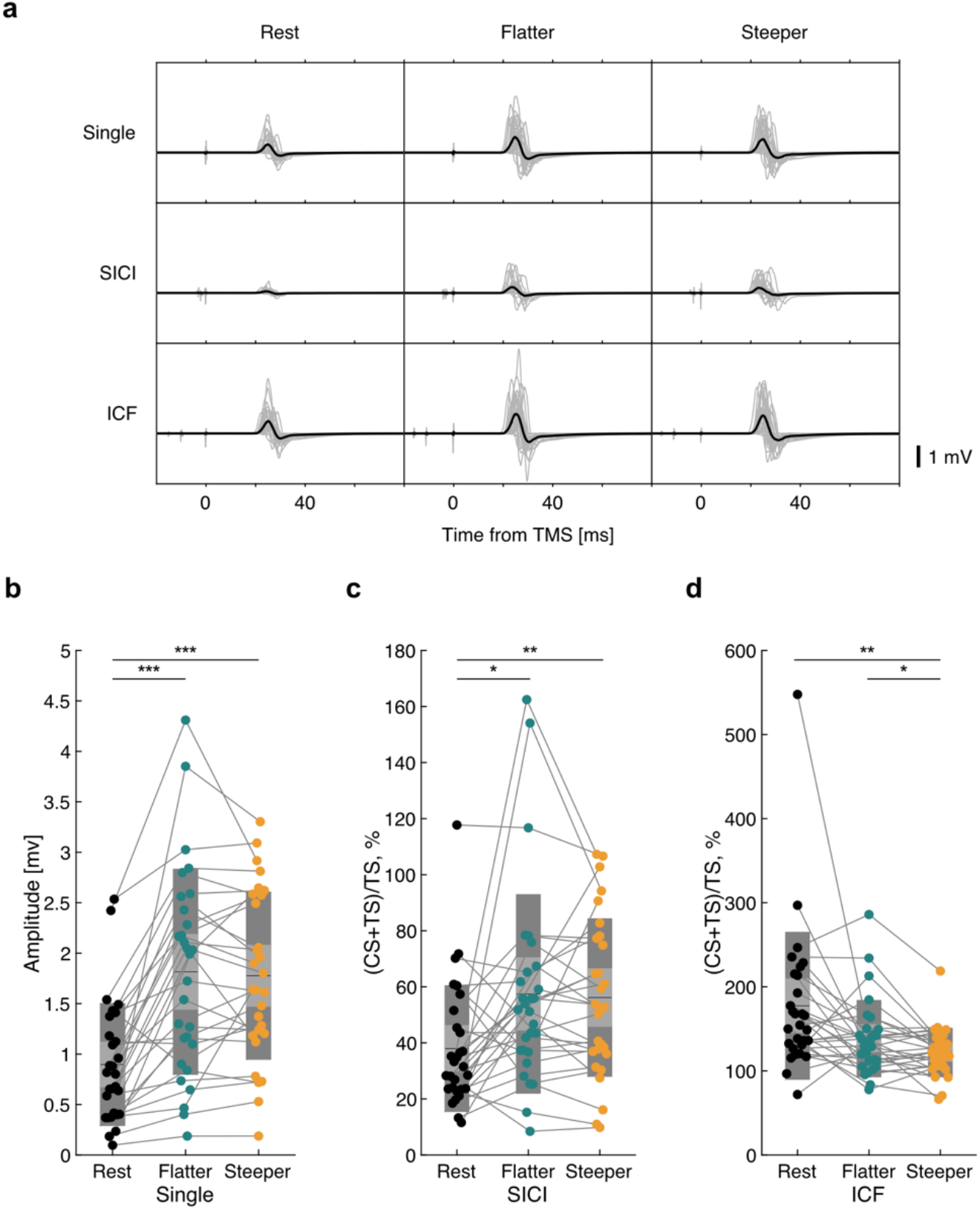
Direction-specific differences in intracortical facilitation across neurofeedback-induced aperiodic states. (a) MEP waveforms elicited by single-pulse TMS, SICI and ICF protocols during the rest, flatter, and steeper states. Gray traces represent the median MEP waveform for each participant within each state and stimulation protocol, and black traces represent the mean of these participant-level median waveforms (n = 29). (b) Single-pulse MEP amplitudes during the rest, flatter, and steeper states. (c) Normalized SICI responses during the rest, flatter, and steeper states. SICI responses were expressed as the conditioned MEP amplitude (CS + TS) relative to the single-pulse MEP amplitude (TS) within each state. (d) Normalized ICF responses during the rest, flatter, and steeper states. ICF responses were expressed as the conditioned MEP amplitude (CS + TS) relative to the single-pulse MEP amplitude (TS) within each state. In panels b–d, dots represent individual participants, gray lines connect measurements from the same participant, and box plots indicate the mean ± 1 SD (dark gray) and the 95% confidence interval of the mean (light gray). Black horizontal lines indicate the group means. Statistical significance is indicated as \**p* < 0.05, \*\**p* < 0.01, and \*\*\**p* < 0.001.

Single-pulse MEP amplitudes showed a neurofeedback-related increase irrespective of modulation direction (Fig. 4b). A one-way repeated-measures ANOVA revealed a significant main effect of state on single-pulse MEP amplitudes (*F*(1.55, 43.53) = 35.04, *p* < 0.001, ηp² = 0.56). Post hoc comparisons showed that MEP amplitudes were significantly greater during both the flatter state (*t*(28) = 6.22, corrected-*p* < 0.001, *d* = 1.10) and the steeper state (*t*(28) = 6.82, corrected-*p* < 0.001, *d* = 1.06) than during the rest state, whereas no significant difference was observed between the flatter and steeper states (*t*(28) = 0.44, corrected-*p* = 1.000, *d* = 0.05), indicating that corticospinal excitability increased during neurofeedback irrespective of modulation direction.

Normalized short-interval intracortical inhibition (SICI) responses showed a similar neurofeedback-related change without a direction-specific difference (Fig. 4c). A one-way repeated-measures ANOVA revealed a significant main effect of state on normalized SICI responses (*F*(1.45, 40.71) = 7.71, *p* = 0.004, ηp² = 0.22). Post hoc comparisons showed that normalized SICI responses were significantly greater during both the flatter state (*t*(28) = 2.78, corrected-*p* = 0.029, *d* = 0.67) and the steeper state (*t*(28) = 3.56, corrected-*p* = 0.004, *d* = 0.63) than during the rest state, whereas no significant difference was observed between the flatter and steeper states (*t*(28) = 0.31, corrected-*p* = 1.000, *d* = 0.05), indicating that normalized SICI responses were similarly greater during both neurofeedback states than during rest, with no direction-specific difference.

In contrast, normalized intracortical facilitation (ICF) responses showed a direction-specific difference between the two neurofeedback conditions (Fig. 4d). A one-way repeated-measures ANOVA revealed a significant effect of state (*F*(1.18, 33.14) = 8.33, *p* = 0.005, ηp^2^ = 0.23). Post hoc comparisons demonstrated significantly lower ICF responses during the steeper state than during the flatter state (*t*(28) = 2.68, corrected-*p* = 0.037, *d* = 0.27). ICF responses during the steeper state were also significantly lower than those during the rest state (*t*(28) = 3.55, corrected-*p* = 0.004, *d* = 0.94), whereas no significant difference was observed between the rest and flatter states (*t*(28) = 2.25, corrected-*p* = 0.098, *d* = 0.66). Thus, unlike single-pulse MEP amplitudes and normalized SICI responses, ICF responses showed a direction-specific difference between the flatter and steeper aperiodic states.

### Trial-wise associations between EEG spectral features and corticospinal and intracortical excitability

The condition-level difference in ICF raised the question of whether prestimulus EEG activity also tracks moment-to-moment fluctuations in motor cortical excitability. At the single-trial level, this relationship could be examined not only for the experimentally targeted aperiodic exponent but also for concurrent fluctuations in periodic spectral activity. We therefore examined trial-wise associations between aperiodic and periodic EEG spectral features and corticospinal and intracortical excitability.

In an initial exploratory screening, trial-wise associations between each EEG feature and each MEP-derived measure were summarized as a heatmap of t values obtained from separate generalized linear mixed-effects models (GLMMs) (Fig. 5a). This screening suggested a negative association between periodic alpha-band power and normalized SICI responses (*t* = −1.92), and a positive association between periodic alpha-band power and single-pulse MEP amplitude (*t* = 1.82). No clear associations involving the aperiodic exponent were apparent for any of the three MEP-derived measures.

**Figure 5.**
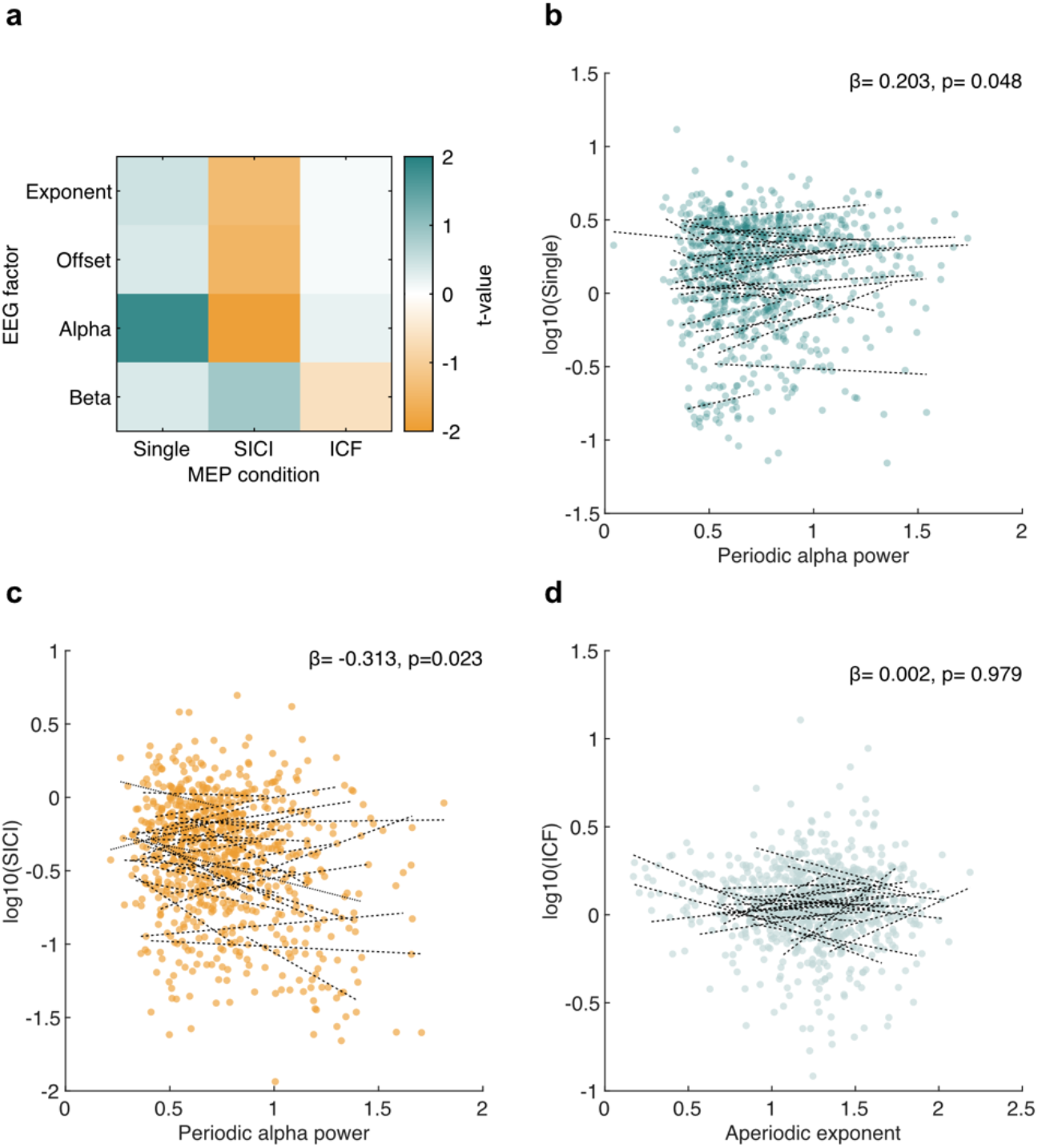
Trial-wise associations between EEG spectral features and corticospinal and intracortical excitability. (a) Exploratory screening of trial-wise associations between prestimulus EEG spectral features and MEP-derived measures. Colors indicate t values obtained from separate GLMMs for each combination of EEG feature and MEP-derived measure. (b) Trial-wise association between periodic alpha-band power and single-pulse MEP amplitude. (c) Trial-wise association between periodic alpha-band power and normalized SICI responses. (d) Trial-wise association between the aperiodic exponent and normalized ICF responses. In panels b–d, dots represent individual trials and black lines represent participant-specific model fits. β coefficients and p values are derived from multivariable GLMMs including the aperiodic exponent, periodic alpha-band power, and periodic beta-band power as predictors. GLMMs were fitted using a Gamma distribution with a log link. MEP-derived measures are shown on a log10 scale for visualization.

We next fitted multivariable GLMMs with the aperiodic exponent, periodic alpha-band power, and periodic beta-band power as predictors. Periodic alpha-band power was positively associated with single-pulse MEP amplitude (β = 0.203, *p* = 0.048; Fig. 5b), whereas neither the aperiodic exponent (β = 0.053, *p* = 0.530) nor periodic beta-band power (β = −0.049, *p* = 0.769) showed significant effects. For normalized SICI responses, periodic alpha-band power showed a significant negative association (β = −0.313, *p* = 0.023; Fig. 5c), indicating that trials with higher prestimulus alpha-band activity tended to exhibit stronger intracortical inhibition. Neither the aperiodic exponent (β = −0.150, *p* = 0.191) nor periodic beta-band power (β = 0.288, *p* = 0.186) significantly predicted SICI responses. For normalized ICF responses, none of the EEG spectral features showed a significant association, including the aperiodic exponent (β = 0.002, *p* = 0.979; Fig. 5d). This lack of a continuous trial-wise association contrasted with the effect of the targeted neurofeedback state itself. ICF was lower in the steeper than in the flatter state (β = −0.10, *p* = 0.002), and this effect was retained when a by-subject random slope for state was included (*p* = 0.017). The steeper state was also characterized by a higher mean aperiodic exponent (Supplementary Fig. 1).

Together, these exploratory analyses indicate that the condition-level difference in ICF was not mirrored by a corresponding trial-by-trial association with the aperiodic exponent. In contrast, periodic alpha-band power showed opposite-signed associations with single-pulse MEP amplitude and normalized SICI responses, although these preliminary associations were modest and uncorrected for multiple comparisons.

## Discussion

The aim of this study was to determine whether volitional modulation of the EEG aperiodic component leads to changes in intracortical excitatory or inhibitory population activity. To this end, we used a closed-loop EEG–TMS protocol to test whether the sensorimotor aperiodic exponent can be brought under voluntary control and whether such control is accompanied by changes in intracortical excitability in human M1. Participants bidirectionally modulated the EEG aperiodic dynamics in the instructed direction within a single session under its real-time feedback, without significant concomitant changes in non-targeted periodic activity (Figs. 2 and 3). Further, TMS-based ICF assessment showed direction-specific modulation (Fig. 4). Together, these findings indicate that brain states characterized by different M1 aperiodic dynamics correspond to distinct intracortical E/I states, suggesting that the neurofeedback-induced aperiodic modulation serves as a macroscopic index of intracortical E/I balance.

Although numerical simulation studies using biophysical models have suggested that synaptic E/I can modulate macroscopic aperiodic neural dynamics^7,9^, its experimental validation for awake healthy humans beyond correlational observations was elusive. To address this, we extended the real-time EEG neurofeedback framework, which to date has primarily targeted periodic features, to the aperiodic component and integrated it with closed-loop brain stimulation. The EEG–TMS paradigms have, to date, been developed almost exclusively around periodic features of EEG, including sensorimotor rhythm power, alpha-band amplitude, event-related desynchronization (ERD), and the phase of the mu rhythm^19–22^. This focus reflects the prevailing view of cortical activity as fundamentally oscillatory^23^.

In parallel, a growing body of work has shown that the aperiodic exponent of the power spectrum is functionally meaningful and tracks physiological and attentional states in a time-resolved manner^10,24,25^. The aperiodic component, however, has primarily been studied as an observational measure rather than a target for active modulation. Whether the closed-loop control established for periodic features can be extended to the aperiodic component itself therefore remained unresolved. Its real-time implementation was also far from trivial, because aperiodic parameterization is sensitive to noise and has conventionally been performed offline. Our findings bridge this gap. By combining short-window *specparam* fitting^4^ with temporal smoothing, we estimated the aperiodic exponent online, and participants were able to shift the spectral slope bidirectionally relative to their own resting state. More broadly, this framework couples neurofeedback-based manipulation of brain state with TMS-based physiological assessment and extends the targets of non-invasive neurofeedback and closed-loop brain stimulation from periodic neural oscillations to broadband aperiodic dynamics.

A notable finding of the present study was that bidirectional neurofeedback modulation of the aperiodic exponent occurred without significant concomitant changes in periodic activity within the theta, alpha, or beta bands (Fig. 3b). This finding suggests that volitional regulation of aperiodic activity does not necessarily require systematic modulation of periodic oscillatory activity. In conventional ERSP, however, significant condition differences were observed in both the theta and beta bands (Fig. 3a). Similar changes across multiple conventional frequency bands have been observed during biofeedback-related modulation^26^. Importantly, changes in the aperiodic component itself can manifest as apparent changes in conventional band-power measures^27^. Thus, the theta- and beta-band differences observed in conventional ERSP were consistent with the spectral consequences expected from the experimentally induced modulation of the aperiodic exponent.

The dissociation between conventional and periodic activity was particularly evident in the theta band. Conventional theta-band ERSP showed a significant difference between the flatter and steeper states, whereas this difference disappeared after removal of the fitted aperiodic component. Although periodic theta activity did not differ significantly between conditions, the numerical direction of the difference was opposite to that observed in conventional ERSP. Similar dissociations between conventional theta-band power and aperiodic-adjusted theta activity have recently been reported, demonstrating that apparent theta-band effects can be substantially altered after accounting for concurrent aperiodic dynamics^28^. Thus, the theta-band difference observed in conventional ERSP was consistent with the spectral consequences of aperiodic modulation rather than reflecting changes in periodic theta-band activity.

A similar pattern was observed in the beta band, where the significant conventional ERSP difference was no longer significant after removal of the aperiodic component. In contrast, alpha activity showed no significant condition difference either before or after aperiodic correction. The absence of a conventional alpha-band difference was also consistent with the induced spectral modulation, as the alpha range was located near the apparent center of the spectral rotation associated with changes in the aperiodic exponent, where the effect of the spectral rotation would be expected to be smaller than the difference at lower and higher frequencies (Fig. 1c). Moreover, the present neurofeedback task did not involve motor imagery or other explicit modulation of sensorimotor rhythms, providing no direct task-related drive for alpha/mu modulation. Together, the absence of significant periodic differences across theta, alpha, and beta band indicates that the experimentally induced bidirectional modulation of the aperiodic exponent was not accompanied by systematic modulation of these oscillatory components.

In computational models, increasing the relative contribution of excitation to inhibition flattens the spectral slope, suggesting that relative changes in E/I balance may in principle be inferred from this spectral feature. This prediction was further supported by a reanalysis of rat hippocampal LFP data, in which the PSD slope covaried with the anatomically defined ratio of excitatory to inhibitory synapse density across CA1 layers^7^. Subsequent biophysical modeling confirmed that the exponent arises from the dynamics of AMPA and GABA_A_ currents, but also showed that it is shaped by several other factors, including leak conductance, and that its relationship to E/I can reverse in sign depending on the conditions. Because a change in the exponent in a given direction can arise from distinct underlying processes, the exponent is not a consistent readout of E/I^9^. All this evidence, however, comes from manipulating E/I and measuring the exponent. Whether a state defined by the exponent itself corresponds to a distinct intracortical state had not been tested.

Our study addresses this gap by directly targeting the aperiodic exponent for experimental manipulation and examining whether the resulting states were accompanied by changes in intracortical physiology. We showed that the aperiodic component was modulated without significant concomitant changes in periodic activity. Under these conditions, ICF was reduced when participants steepened the exponent (Fig. 4d). ICF is influenced by both facilitatory and inhibitory processes within intracortical circuits rather than selectively reflecting a specific neurotransmission mechanism, since ICF depends on NMDA receptor-mediated glutamatergic transmission and is also reduced by positive modulators of GABA_A_ receptors^18,29–31^. Notably, whether the reduction in ICF arises from decreased facilitation or increased inhibition, either would be consistent with a shift toward lower E/I, as theoretically predicted for a steeper aperiodic exponent.

This interpretation is supported by the absence of a difference in single-pulse MEP amplitude between the flatter and steeper states (Fig. 4b), which indicates that the state difference in ICF does not depend on a difference in the magnitude of the test-stimulus response. Single-pulse MEP amplitude was nevertheless larger in both feedback conditions than at rest. Because a pre-recorded feedback was displayed during the resting condition to match visual input, this increase is unlikely to reflect visual stimulation itself, and more likely reflects non-specific differences between actively performing the task and being at rest, such as attention, arousal, and task engagement. Corticospinal excitability is indeed known to vary with visual attention and arousal state^12,32^. Critically, because this non-specific increase was common to both feedback conditions, it does not affect the comparison between the flatter and steeper conditions.

At first sight, the finding that SICI, which indexes the strength of GABA_A_ergic circuits^16,18^, did not differ between the flatter and steeper states appears inconsistent with a change in E/I (Fig. 4c). However, this can be explained by the fact that modulation of the EEG aperiodic component was achieved here without accompanying modulation of the periodic component. Previous work has shown that ERD is accompanied by a reduction in SICI but leaves ICF unaffected^33^, suggesting a closer relationship between SICI and periodic alpha-band activity. Consistent with this, in our trial-level analysis, the normalized SICI response was related to periodic alpha-band activity (Fig. 5c). The invariance of SICI across the two neurofeedback states, in which the periodic component did not change, is therefore consistent with the previous finding and does not contradict our ICF-based interpretation. A comparable dissociation has been reported previously. The N45 component of the TMS-evoked potential, which is thought to reflect GABA_A_ receptor-mediated neurotransmission^34^, does not correlate with the aperiodic exponent within the same participants, and the two differ in both their direction of change and their relation to behavior, suggesting that the N45 component and the aperiodic exponent capture different aspects of inhibition^35^.

Our finding that modulation of the aperiodic component occurred without a change in SICI is consistent with this dissociation. SICI was nonetheless weaker in both neurofeedback conditions relative to rest (Fig. 4c). This is consistent with the non-specific effects of task engagement noted above, since visual attention has been reported to weaken SICI while leaving ICF unchanged^32^. Attentional engagement during the task may therefore have contributed to the difference in SICI from rest, whereas the same factor cannot account for the ICF difference between the flatter and steeper states (Fig. 4d).

Although the difference in ICF between the flatter and steeper states suggests an association between aperiodic activity and intracortical physiology at the state level (Fig. 4d), a significant association was not observed at the trial level. Specifically, the continuous trial-by-trial aperiodic exponent was not significantly associated with ICF in a multivariable model that also included periodic alpha- and beta-band power (Fig. 5d). Thus, the observed state-dependent difference in ICF did not reflect a one-to-one correspondence between the aperiodic exponent value and ICF on individual trials. This dissociation between the state- and trial-level associations is consistent with the notion that the aperiodic exponent is not uniquely determined by a single physiological process. Different cellular, synaptic, circuit, and measurement-level processes can generate similar finite-band spectral exponents, limiting mechanistic identifiability from the exponent alone^13^. Consistent with this principle, biophysical models have demonstrated that similar aperiodic exponent values can arise from different combinations of synaptic currents and membrane properties^9^. Thus, comparable exponent values do not necessarily indicate the same intracortical physiological state. Trial-to-trial fluctuations in the aperiodic exponent may therefore have contained not only components related to the net intracortical E/I state reflected by ICF, but also physiological variability that is not captured by ICF. Together, these findings support the interpretation that the aperiodic exponent reflects the intracortical E/I state captured by ICF at the macroscopic level, rather than providing a direct readout of the underlying E/I balance.

Several limitations should be considered. First, our final sample reflects considerable attrition. Of 47 volunteers, 17 were excluded because their resting motor threshold (rMT) exceeded 84% of maximum stimulator output, and of the 30 participants who completed the protocol, one was further excluded because of physiologically implausible aperiodic estimates, leaving 29 for analysis. This attrition follows directly from the stringent TMS and EEG quality-control criteria that we imposed. Most exclusions arose because, in participants with a high rMT, the suprathreshold test-stimulus intensity required for paired-pulse TMS could not be delivered within the output range of the stimulator. In addition, reliable estimation of the aperiodic exponent requires spectra of sufficient quality and careful parameterization^4,36^, and brain-state-dependent TMS based on real-time EEG has required selecting participants in whom the targeted neural activity can be clearly detected^21^. We therefore regard this attrition not as a measurement shortcoming but as the cost of acquiring and stimulating a genuinely reliable physiological target. It nonetheless limits the generalizability of our findings and may bias the sample towards individuals with favourable corticospinal excitability and signal quality. Second, voluntary aperiodic modulation cannot be fully dissociated from arousal-related changes accompanying task engagement. Task engagement flattens the spectrum in the cortical regions involved^10,37^, and increased arousal, as indexed by pupil size, has been associated with a flatter spectral slope^12^. The flatter state may therefore have included flattening driven by attention and arousal rather than by volitional modulation alone. Critically, visual attention has been reported to leave ICF unchanged despite increasing corticospinal excitability and reducing SICI^32^. If the exponent reduction in the flatter condition arose largely from attention-related flattening, the absence of an ICF difference between the rest and flatter conditions (Fig. 4d) is what would be predicted. Designs that manipulate arousal independently of the aperiodic target, or that require flattening beyond the level reached by ordinary task concentration, would allow this confound to be disentangled and would test whether truly symmetric bidirectional control is attainable.

In summary, neurofeedback enabled participants to bidirectionally modulate the aperiodic component without significant concomitant changes in periodic activity. This manipulation was accompanied by a direction-specific change in ICF that was consistent with E/I-based predictions, supporting the interpretation that the aperiodic exponent reflects, at the macroscopic level, the intracortical E/I state captured by ICF rather than providing a direct readout of the underlying E/I balance. More broadly, our findings suggest that the aperiodic component may serve not only as a marker of cortical physiological state, but also as an intervention target through which E/I-related intracortical physiology can be modulated in a desired direction. Whereas previous studies have altered E/I states through pharmacological interventions or exogenous stimulation, our findings suggest that voluntary modulation of the aperiodic component through neurofeedback may provide a means of inducing distinct states of intracortical excitability. Given that E/I balance predicts subsequent learning and responsiveness to neural stimulation^38,39^, adjusting the pre-intervention intracortical state in a desired direction through aperiodic neurofeedback may provide a new approach for optimizing the induction of neuroplasticity according to the individual’s state.

## Methods

### Study design

This randomized within-participant study tested whether bidirectional neurofeedback targeting the sensorimotor EEG aperiodic exponent could modulate corticospinal and intracortical excitability in the contralateral M1. Each participant completed three experimental conditions: a resting-state baseline condition and two neurofeedback conditions. Reducing and increasing aperiodic exponent (i.e., flatter and steeper power spectra) were required in the flatter and steeper neurofeedback conditions, respectively. The order of the two neurofeedback conditions was randomized across participants to control for order effects.

Before neurofeedback, resting-state EEG recordings and TMS measurements were acquired. The resting-state TMS measurements were used both to determine individualized interstimulus intervals (ISIs) for paired-pulse TMS and to provide resting-state measures of single-pulse MEP amplitude, SICI, and ICF. During the neurofeedback conditions, participants received real-time feedback designed to bias the aperiodic exponent of contralateral M1 sensorimotor activity toward either a flatter or steeper state.

The target sample size was determined a priori on the basis of previous within-participant studies that combined EEG with single-pulse or paired-pulse TMS to relate sensorimotor brain states to corticospinal and intracortical excitability^22,33,40,41^, in which 20 to 31 participants were tested. We therefore set a target of 30 participants completing the experimental protocol.

### Participants

Forty-seven volunteers (13 women and 34 men; mean age ± SD, 21.9 ± 1.93 years; range, 19–30 years) were recruited for this study. All participants had normal or corrected-to-normal vision and reported no history of neurological or psychiatric disorders. Handedness was assessed using the Edinburgh Handedness Inventory, and all participants were right-handed (laterality quotient, 88.1 ± 12.4 %).

Seventeen participants were excluded from the TMS experiment because the resting motor threshold of the right first dorsal interosseous muscle exceeded 84% of the maximum stimulator output. This exclusion criterion was applied to ensure that the stimulator could deliver the required stimulation intensities throughout the experiment. A relatively large portion of participants were excluded because the EEG cap increased the resting motor threshold^42^.

Thirty participants completed the experimental protocol. One participant was subsequently excluded from all EEG-triggered TMS analyses due to compromised signal quality. Thus, the final sample consisted of 29 participants (8 women and 21 men; mean age ± SD, 21.8 ± 2.08 years; range, 19–30 years). Among the 29 participants included in the analyses, fifteen started with the flatter-feedback condition and fourteen started with the steeper-feedback condition.

All procedures conformed to the Declaration of Helsinki and the current safety guidelines for TMS issued by the International Federation of Clinical Neurophysiology. The experimental protocol was approved by the Ethics Committee of the Faculty of Science and Technology, Keio University (approval no. 2025-007). All participants provided written informed consent before participation.

### EEG/EMG data acquisition

Scalp EEG was recorded using a 64-channel whole-head EEG sensor net (MicroCel Geodesic Sensor Net, Electrical Geodesics Inc., Eugene, OR, USA). Conductive EEG paste (Elefix V ZV-181E, Nihon Kohden, Tokyo, Japan) was applied between the scalp and electrodes to reduce electrode impedance. EEG data quality was maintained for channels used in the primary analysis over the sensorimotor cortex, with impedances kept below 50 kΩ. Some non-primary channels could not be fully reduced due to individual differences in head size and hair volume, but these were not included in the main analysis.

Surface EMG was recorded from the first dorsal interosseous muscle. Disposable surface electrodes with a diameter of 24 mm (Spes Medica, Genova, Italy) were attached over the muscle. Before data acquisition, EMG signals were visually inspected to confirm the absence of environmental noise and voluntary muscle activity, and the baseline offset was corrected.

### TMS-Robot and RGBD-based neuronavigation

Corticospinal excitability was assessed using a robotic TMS system^43^. Magnetic pulses were delivered through an actively cooled figure-of-eight coil (Cool-B65 coil; MagVenture A/S) connected to a TMS stimulator (MagPro X-100; MagVenture A/S, Farum, Denmark) and the coil was mounted on a six-axis collaborative robotic arm. The robot maintained the coil tangential to the scalp at the predefined stimulation target, while the position of the participant’s head and the coil was tracked in real time using an optical tracking system. Stimulation was automatically paused when the coil deviated from the target position or orientation beyond the predefined tolerance.

For MRI-free neuronavigation, an individual three-dimensional head model was generated using an RGBD depth camera. Participants wore a tight-fitting cap to reduce hair-related surface artifacts, and the scalp and facial surfaces were scanned before TMS. The reconstructed mesh was inspected for visible defects and imported into the robotic TMS system. Head-to-model registration was performed using anatomical landmarks, including the nasion, inion, and bilateral preauricular points. Registration was accepted only when the residual landmark error was within 5 mm. Additional scalp surface points were then sampled and used to refine the registration by an iterative closest-point algorithm.

The resting motor threshold was estimated at an initial stimulation site around the left primary motor cortex representation of the target hand muscle. After threshold estimation, motor mapping was performed at 110% of the resting motor threshold. Stimulation sites were sampled around the initial target, and MEPs were recorded from the target hand muscle. The motor hotspot was defined as the center of gravity of the motor map, computed by weighting each stimulation coordinate by the corresponding peak-to-peak MEP amplitude. The resting motor threshold was then re-estimated at this hotspot and used for the subsequent experimental stimulation protocol.

### ISI evaluation

Before the main experiment, we determined ISI for paired-pulse TMS on an individual basis. Because the optimal ISI for paired-pulse TMS varies across individuals, we evaluated four commonly used ISIs: 2 and 3 ms for SICI, and 10 and 15 ms for ICF. Five stimulation conditions were tested: single-pulse TMS and paired-pulse TMS with ISIs of 2, 3, 10, and 15 ms. Each condition was delivered 10 times in a randomized order while participants remained at rest. The interval between consecutive stimulation trials was 4.0 ± 0.5 s.

During this evaluation, participants viewed the same neurofeedback display as that used in the flatter-feedback, and steeper-feedback conditions of the main experiment, but were instructed to remain relaxed and passively observe the display without attempting to enlarge the feedback circle. Thus, the visual input was matched across the ISI evaluation and the main experimental conditions, allowing the MEPs obtained during this procedure to be treated as resting-state MEP data.

### Experimental protocol and aperiodic neurofeedback

Each trial consisted of four phases: rest, ready, task, and blank (Supplementary Fig. 2). During the rest phase, no visual feedback was provided, and the baseline aperiodic exponent was estimated from the 5-s resting EEG segment.

Participants were instructed to remain still and avoid blinking, swallowing, and voluntary muscle contraction to minimize non-neural EEG and EMG artifacts. During the rest phase, participants were instructed to relax and look at the feedback display. The ready phase served as a preparatory cue indicating that the task phase would begin shortly. During the task phase, participants were instructed to try to enlarge and maintain the feedback circle. During the blank phase, participants were allowed to relax, and small movements such as blinking were permitted before the next trial.

The task-related modulation of the aperiodic component was quantified as:

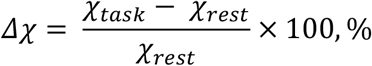

where ***χ_task_*** and ***χ_rest_*** denote the aperiodic exponent during the task and rest phases, respectively. Because ***χ_task_*** was updated every 100 ms, Δχ was also updated every 100 ms and used for real-time visual feedback.

We presented visual feedback of the real-time calculated Δχ value as a purple circle on a computer screen, the size of which reflected the magnitude of Δχ in the instructed target direction, with Δχ = 0 % as the baseline and 15 % as maximum. Participants were not informed of the target direction or its changes between blocks. Participants were instructed only to make the purple circle appear, enlarge it, and maintain its size when it reached the maximum (i.e., keeping 15% modulation).

Neurofeedback trials were organized into blocks of 15 trials. The experiment continued until 12 triggered MEPs were obtained for each stimulation type—single-pulse, SICI, and ICF—in each neurofeedback condition. The number of blocks therefore varied across participants depending on their ability to modulate the aperiodic exponent and ranged from 6 to 12 blocks. The protocol was designed to terminate after 12 blocks even if the target number of triggered trials had not been reached. However, all participants reached the required number of triggered MEPs within 12 blocks.

### Adaptive randomization of TMS conditions

Within the first block, the three TMS conditions were initially balanced, with 5 trials each assigned to single-pulse TMS, SICI, and ICF, yielding 15 TMS trials in total. The order of stimulation conditions was randomized within the block.

From the second block onward, the allocation of stimulation conditions was adaptively adjusted according to the number of successfully triggered trials obtained for each stimulation condition in the preceding blocks. Specifically, the number of successful triggered trials was counted separately for single-pulse TMS, SICI, and ICF. Stimulation conditions with fewer accumulated successful trials were assigned a higher probability of being selected in the next block, whereas conditions with more accumulated successful trials were assigned a lower probability. This procedure was used to balance the final number of successfully triggered MEPs across the three stimulation conditions.

For each subsequent block, the selection weight for each stimulation condition was calculated as:

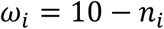

where n_i_ denotes the accumulated number of successfully triggered trials for stimulation condition i. If n_i_ exceeded 10, it was truncated to 10. The probability of assigning stimulation condition i was then calculated as:

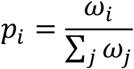

The expected number of trials for each stimulation condition in the next block was calculated by multiplying p_i_ by the total number of TMS trials in the block. These expected values were converted to integer trial counts by taking the floor value, and any remaining trials were assigned to conditions with the largest fractional remainders. The resulting labels were randomly permuted to determine the stimulation order within the block.

### Online EEG processing and neurofeedback

EEG signals were recorded at 1,000 Hz and downsampled to 200 Hz for online analysis. Five channels over the left sensorimotor cortex were selected as channels of interest and re-referenced based on the Laplacian montage^44^.

Raw EEG data were analyzed every 100 ms. At each update, the signal was band-pass filtered between 1 and 70 Hz and notch-filtered at 50 Hz. A small Laplacian spatial filter centered on C3 was applied by subtracting the mean of the four surrounding electrodes from the center electrode.

Spectral power was estimated using fast Fourier transform (FFT) applied to a 2-s EEG segment spanning from 2.2 to 0.2 s before each update. This 200-ms analysis offset was introduced to reduce filter-edge artifacts. The analysis was repeated every 100 ms, and PSD estimates from the most recent 10 analyses were averaged to reduce short-term spectral fluctuations. This procedure prioritized stable online estimation over instantaneous feedback. Therefore, the feedback signal reflected temporally smoothed aperiodic activity rather than a single-segment estimate.

The aperiodic exponent of the power spectrum was estimated using *specparam* algorithm. Frequencies were defined from 0.5 to 40 Hz in 0.5-Hz increments, and model fitting was performed over the 2–40 Hz range. The parameterization algorithm was configured with a fixed aperiodic mode, peak width limits of 0.5–12.0 Hz, no upper limit on the number of peaks, a minimum peak height of 0.0, and a peak threshold of 2.0. The same configuration was used for all subsequent analysis. Negative aperiodic exponent estimates were treated as invalid in the present fitting range and preprocessing pipeline, because they likely reflected unstable model fitting rather than physiologically interpretable 1/f-like spectral structure. This criterion was applied to all analyses reported below.

At the beginning of each ready phase, a reference aperiodic exponent was calculated from the preceding baseline samples. Because online analysis was performed every 100 ms over a 5-s period, up to 50 samples were available for reference estimation. Successfully parameterized samples were further screened for outliers and those outside mean ± 3SD were excluded from further analysis. During the task phase, the aperiodic exponent was updated every 100 ms and converted into the percentage modulation Δχ defined above, with this reference value serving as the rest-phase exponent.

### Brain-state-dependent TMS triggering

Threshold monitoring began 1 s after task onset, corresponding to 7 s after trial onset. TMS was triggered when the feedback signal reached or exceeded the target threshold of 15% in the instructed direction (Supplementary Movie 1).

A fallback trigger time was randomly selected within the 11–12 s interval after trial onset in 100-ms increments. If the feedback signal reached the target threshold before this fallback time, TMS was delivered immediately. Otherwise, TMS was delivered at the fallback time, ensuring that one TMS stimulation event was administered in each trial.

### MEP analysis

MEP analysis was performed offline using custom MATLAB scripts. Trials were excluded from analysis according to predefined quality-control criteria based on EMG activity, TMS coil positioning, and stimulation validity.

First, trials were excluded when reliable stimulation delivery could not be ensured. Specifically, trials were manually marked as “Rejected” by the experimenter when excessive participant movement, voluntary muscle contraction, or inappropriate coil positioning was observed at the time of stimulation. In addition, trials were excluded when the coil pressure fell below 0.1, when the angular error exceeded 5°, or when the stimulation position deviated by more than 5 mm from the predefined hotspot.

Second, trials with insufficient EMG responses were excluded using a baseline-dependent threshold. Baseline EMG activity was evaluated during the prestimulus period from −200 to 0 ms relative to the TMS pulse. For each trial, the standard deviation of the baseline EMG was calculated, and trials were rejected when the maximum absolute poststimulus EMG amplitude did not exceed five times the baseline standard deviation. This procedure was used to exclude trials in which a reliable MEP response could not be identified.

In addition, only brain-state-triggered trials were included in subsequent analyses. Trials in which stimulation was delivered at the fallback trigger time (non-trigger trials) were excluded from both EEG and MEP analyses because stimulation timing in these trials was not contingent on the target brain state. MEPs obtained during the resting baseline, flatter-feedback, and steeper-feedback conditions were classified into three stimulation conditions. Single-pulse TMS, SICI and ICF. Peak-to-peak MEP amplitude was defined as the difference between the maximum and minimum EMG amplitudes within the 20–60 ms poststimulus window. For each participant and condition, mean peak-to-peak MEP amplitudes were calculated separately for single-pulse TMS, SICI, and ICF trials. The mean SICI and ICF MEP amplitudes were then divided by the corresponding mean single-pulse MEP amplitude obtained from the same participant and condition. Thus, conditioned responses were expressed as ratios relative to the corresponding single-pulse MEP amplitude.

### Offline EEG preprocessing and spectral analysis

Offline preprocessing was designed to estimate the pre-stimulation cortical state using procedures conceptually similar, but not identical, to the online neurofeedback pipeline. To evaluate the pre-stimulation primary motor cortical state that may influence corticospinal excitability, offline EEG preprocessing and spectral analyses were performed.

The time segment of interest was defined as the pre-stimulation period preceding the TMS pulse, in order to avoid contamination by TMS-induced artifacts. Continuous EEG signals were first inspected for TMS pulse artifacts. TMS artifacts were detected using an amplitude-based peak detection procedure, and EEG samples from 50 ms before to 500 ms after each TMS pulse were marked as artifact-contaminated. These samples were temporarily replaced with NaN values and linearly interpolated along the temporal dimension. The artifact mask was retained for subsequent reapplication after filtering procedures.

The same small Laplacian spatial filter as in the online analysis was then applied to enhance local sensorimotor activity, subtracting the average activity of the surrounding electrodes from the center electrode.

EEG data were subsequently epoched relative to experimentally recorded stimulation and rest timestamps. Stimulation epochs ranged from −5 to +2 s relative to TMS onset, whereas resting epochs consisted of 5-s segments extracted from the resting period. Epoched EEG signals were band-pass filtered between 1 and 70 Hz and notch-filtered at 50 Hz using zero-phase forward and reverse filtering.

After filtering, artifact-contaminated samples identified by the retained artifact mask were set to zero again to minimize residual interpolation-related contamination in subsequent spectral analyses.

For spectral analysis, the channel of interest corresponding to the left primary motor cortex was extracted. Time-resolved power PSD estimates were calculated using a sliding-window FFT with a 1-s Hanning window and 90% overlap between consecutive windows, corresponding to a 100-ms temporal update interval. To reduce edge-related spectral artifacts, EEG epochs were zero-padded before FFT analysis. The resulting PSD estimates were subsequently smoothed using a moving average over the current and preceding 10 spectral windows while omitting missing values.

The pre-stimulation spectral segment was extracted from the smoothed PSD estimates immediately before TMS onset. Specifically, the final 20 spectral windows up to the TMS trigger window were selected and used for spectral parameterization. For resting epochs, smoothed PSD estimates from the resting period were used as baseline spectral estimates.

The aperiodic exponent of the EEG power spectrum was parameterized using *specparam* with the configuration described above, with spectral estimates obtained at 1-Hz resolution and model fitting performed over the 2–40 Hz range. These parameters allowed narrowband oscillatory peaks to be modeled separately from the broadband aperiodic component.

For each trial and time window, the aperiodic exponent was extracted from the fitted *specparam* model, and negative estimates were replaced with NaN. For the flatter- and steeper-feedback conditions, the pre-stimulation aperiodic exponent was calculated as the mean of the valid estimates across the 20 spectral windows immediately preceding TMS onset. Trial-wise pre-stimulation exponent values were then screened for outliers separately for each participant and feedback condition using a median-based outlier detection method, and identified outliers were replaced with NaN.

For the resting condition, the aperiodic exponent was estimated using the same preprocessing and spectral parameterization procedures. Resting EEG obtained during the ISI evaluation procedure was not used for spectral estimation because TMS pulses were delivered every 4.0 ± 0.5 s, resulting in substantial contamination of the continuous EEG signal by TMS-related artifacts. Therefore, the final resting value was calculated as the mean of valid estimates across 600 spectral windows obtained from a dedicated 1-min eyes-open resting EEG recording.

### Conventional periodic ERSP analysis

To determine whether neurofeedback-induced modulation of the aperiodic exponent was accompanied by changes in sensorimotor oscillatory activity, ERSP was quantified in the theta, alpha, and beta bands. All band-power calculations were performed on the C3 Laplacian-filtered EEG signal described above. The theta and beta bands were defined as 4–8 Hz and 13–30 Hz, respectively. For alpha, an individual alpha frequency (IAF) approach was used. IAF was estimated from the dedicated 1-min eyes-open resting-state EEG recording using *specparam* parameterization of the averaged resting power spectrum. The IAF was defined as the frequency of the largest periodic peak within the 7–14 Hz range. When multiple periodic peaks were identified within this range, the peak exhibiting the greatest power was selected. The estimated IAF was rounded to the nearest integer frequency, and an individualized alpha band was defined as IAF ± 1 Hz. For two participants, no periodic peak was identified within the 7–14 Hz range in the C3 Laplacian-filtered signal; in these cases the IAF was estimated from the homologous right sensorimotor channel (C4), and the resulting individualized alpha band was subsequently applied to all analyses for that participant. For each trial, theta-, alpha-, and beta-band power was calculated from the same pre-stimulation EEG segment used for aperiodic exponent estimation. Baseline power was calculated from the corresponding rest period within the same trial using the identical frequency bands. ERSP was computed as the percentage change in band power relative to the trial-specific baseline:

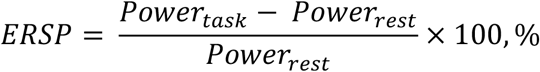

Trial-wise ERSP values were averaged within each participant and neurofeedback state after removal of outliers identified by a median-based criterion, yielding one value per participant, state, and frequency band.

### Cluster-based permutation testing of aperiodic ERSP

To assess changes in aperiodic power across time and frequency, aperiodic power was reconstructed for each time window from the offset and exponent estimated using the *specparam* algorithm as power = offset − χ × log10(f), where χ represents the aperiodic exponent and f represents frequency. Aperiodic power was reconstructed across 2–40 Hz in 1-Hz steps. Aperiodic ERSP was then calculated relative to resting-state aperiodic power.

To determine whether aperiodic ERSP differed from the resting-state reference across time and frequency, cluster-based permutation tests were performed separately for the Flatter and Steeper conditions using FieldTrip. For each condition, participant-level aperiodic ERSP maps were tested against zero using a two-sided dependent-samples t-statistic within a Monte Carlo framework. Because an aperiodic ERSP value of zero indicated no change from rest, this analysis tested for significant increases or decreases in aperiodic power relative to the resting-state reference. Time–frequency bins exceeding a cluster-forming threshold of p < 0.05 were grouped into contiguous clusters across time and frequency. Cluster-level statistics were calculated using the maxsum method, in which t values were summed within each cluster. Statistical significance was assessed using 5,000 randomizations, with pcluster < 0.05 considered significant. Positive and negative clusters were evaluated within the two-sided test. Reported temporal and spectral boundaries represent the observed extent of significant clusters and do not imply pointwise significance at every time–frequency bin within those boundaries.

### Cortical source estimation and parcel-wise aperiodic exponent

To determine whether the condition-dependent change in the aperiodic exponent was present in the cortex underlying the electrode used for neurofeedback, the pre-stimulation EEG was projected into source space. This analysis used a separate preprocessing pipeline from the C3 Laplacian analysis described above. Because the small Laplacian spatial filter is itself a spatial high-pass filter that largely suppresses artifacts arising outside the sensorimotor region, whereas source reconstruction uses all scalp channels and is therefore sensitive to artifacts on any of them, artifact removal by independent component analysis and bad-channel handling were applied only to the data used for source reconstruction.

Continuous EEG was first submitted to an automated preprocessing pipeline implemented in EEGLAB. Channel quality was assessed by fitting the power spectrum of each channel with *specparam*; channels whose aperiodic exponent was negative, or for which the model failed to converge, were marked as bad (0.69 ± 1.01 channels per block, mean ± SD). Line noise at 50 Hz was removed with CleanLine, and amplitude outliers were identified from the mean absolute amplitude of consecutive 0.25-s segments using a median-based criterion. Artifact Subspace Reconstruction was then applied with a cutoff of 5, calibrated on the first 2 s of each epoch (−5.0 to −3.0 s relative to the TMS pulse), which did not overlap with the analysis window defined below. Together with the channels identified at this stage, a total of 5.55 ± 1.34 channels per block were marked as bad. Independent component analysis (binica) was performed on the remaining channels, the channels marked as bad were restored by spherical spline interpolation, and components classified as non-neural by ICLabel were removed (12.2 ± 3.8 of 58.3 ± 1.9 components, 21%).

Cortical sources were reconstructed with sLORETA in Brainstorm. The source space was the cortical surface of the ICBM152 template anatomy (15,002 vertices), and the forward model was computed with a three-shell boundary element method (OpenMEEG; scalp, skull and brain conductivities of 0.33, 0.0042 and 0.33 S/m). Dipole orientation was constrained to the cortical surface normal, the noise covariance regularization parameter was set to 0.1, and the assumed signal-to-noise ratio was 3. Vertex time series were summarized into the 100 parcels of the Schaefer atlas (7-network version); the representative time series of each parcel was the first principal component of its constituent vertex time series. All subsequent analyses were performed on the 100 parcel time series, resampled to 250 Hz.

For each trial, time-resolved PSD estimates were computed with the same sliding-window FFT as in the sensor-space analysis but without zero-padding, yielding 41 spectral windows per 5-s epoch. The final 20 spectral windows up to the TMS trigger were selected and averaged, and the averaged spectrum was used for spectral parameterization (spanning 2.9 s; −2.904 to −0.008 s relative to the TMS pulse). This interval matches the effective integration window of the online feedback signal, which averaged the ten most recent estimates of a 2-s segment updated every 100 ms and therefore reflected 2.9 s of EEG (−3.1 to −0.2 s relative to each update). The aperiodic exponent was parameterized using *specparam* over the 2–40 Hz range at 1-Hz resolution, with the same configuration as the sensor-space analysis. No parcel yielded a negative aperiodic exponent. For each participant and condition, the representative value of each parcel was the median across trials. The cortical region corresponding to the neurofeedback electrode was defined a priori as the region of interest.

The cortical vertex nearest to electrode C3 (15.0 mm from the electrode; MNI −49, −17, 61) fell within the left somatomotor parcel 5 of the Schaefer atlas (7Networks_LH_SomMot_5), which straddles the central sulcus and covers the hand region of the left pre- and postcentral gyri. Its centroid (MNI −40, −23, 57) lay 2.7 mm from the canonical hand knob of the left primary motor cortex (MNI −38, −22, 56). This parcel was defined as the cortical region corresponding to the targeted C3 electrode.

For each of the 100 cortical parcels, the aperiodic exponent was compared between the flatter-feedback and steeper-feedback conditions using a paired-samples t test. The condition difference was defined as flatter-feedback minus steeper-feedback, such that negative values indicated a larger aperiodic exponent in the steeper-feedback condition. No values were missing, yielding 28 degrees of freedom for all parcel-wise comparisons. To account for multiple comparisons across the 100 parcels, p values were corrected using the Benjamini–Hochberg false discovery rate procedure, with FDR-corrected p < 0.05 considered statistically significant. FDR-corrected values are denoted p_FDR. Cohen’s d_z_ was calculated to quantify the within-participant effect size.

### Trial-by-trial associations between prestimulus EEG features and MEP-derived measures

To explore whether trial-by-trial fluctuations in prestimulus EEG activity were related to corticospinal and intracortical excitability, we performed secondary exploratory analyses relating EEG-derived spectral features to MEP-derived measures on a per-trial basis.

EEG features were obtained from the same C3 Laplacian pre-stimulation spectra used for the aperiodic exponent analysis described above. Four features were extracted for each trial: the aperiodic exponent, the aperiodic offset, periodic alpha-band power, and periodic beta-band power. Periodic power was defined as the height of the largest periodic peak detected within the corresponding frequency band, that is, the elevation of the peak above the aperiodic component in log10 power units (the peak power parameter of the *specparam* model). Fixed frequency bands were used, 8–13 Hz for the alpha band and 13–30 Hz for the beta band. When no periodic peak was detected within a band, the feature was treated as missing for that trial.

Three MEP-derived measures were used: single-pulse MEP amplitude, normalized SICI responses, and normalized ICF responses. For SICI and ICF, the conditioned MEP amplitude of each trial was divided by the participant’s mean single-pulse MEP amplitude obtained within the corresponding feedback condition, yielding trial-wise indices of intracortical inhibition and facilitation. This trial-wise normalization differs from the participant-level ratios used for the primary analyses, in which condition means were computed before division.

Only successfully triggered trials were included. Trials whose pre-stimulation aperiodic exponent had been identified as an outlier by the median-based criterion described above were excluded before model fitting, and trials with any missing EEG feature were excluded listwise. This left 29 participants and 793, 751, and 767 trials for the single-pulse, SICI, and ICF models, respectively.

All models were GLMMs with a Gamma distribution and a log link, fitted by maximum pseudo-likelihood (fitglme, MATLAB R2024b; MathWorks, Natick, MA). Fixed effects were tested using Wald t statistics with residual degrees of freedom.

As an initial exploratory screening, 12 separate models were fitted, one for each combination of the four EEG features and the three MEP-derived measures. Each model included a single EEG feature as a fixed-effect predictor and a participant-specific random intercept, and the resulting t statistics were summarized as a heatmap. Because this step was intended solely as a screening procedure, no correction for multiple comparisons was applied and the t statistics are reported descriptively.

To further characterize the associations suggested by this screening, multivariable models were fitted separately for each MEP-derived measure, with the aperiodic exponent, periodic alpha-band power, and periodic beta-band power entered simultaneously as fixed effects together with a participant-specific random intercept. The aperiodic offset was not included because it was strongly collinear with the aperiodic exponent (r = 0.80–0.86 across MEP conditions). To compare the trial-wise effect of the neurofeedback state with the EEG-feature associations, an additional GLMM was fitted to normalized ICF responses with state as a fixed effect and a participant-specific random intercept. Robustness was assessed by refitting the model with an additional by-participant random slope for state.

### Statistical analysis

Statistical analyses were performed using JASP (version 0.95.4). Statistical significance was set at α = 0.05, and all tests were two-sided unless otherwise specified. For comparisons involving three or more repeated-measures conditions, one-way repeated-measures analyses of variance (ANOVAs) were performed. Sphericity was assessed using Mauchly’s test, and Greenhouse–Geisser correction was applied when the assumption of sphericity was violated. When a significant main effect was identified, pairwise comparisons were performed using paired-samples *t*-tests with Bonferroni correction for multiple comparisons.

Time–frequency differences in aperiodic ERSP relative to rest were assessed using cluster-based permutation tests as described above. Statistical significance was determined at the cluster level at *p*_cluster_ < 0.05.

For source-space analyses, the aperiodic exponent was compared between the flatter and steeper conditions within each of 100 cortical parcels using paired-samples *t*-tests. Multiple comparisons across parcels were controlled using the Benjamini– Hochberg FDR procedure, with an FDR-corrected threshold of *p*_FDR_ < 0.05. Effect sizes are reported as partial eta squared (ηp²) for repeated-measures ANOVAs, Cohen’s *d*_z_ for paired-samples *t*-tests, and Cohen’s *d* for post hoc pairwise comparisons.

## Acknowledgements

This work was supported by JST, PRESTO Grant Number JPMJPR23I1, Japan and JST Moonshot R&D Grant Number JPMJMS2012, Japan. The authors thank MEDICAL FIG., a service of Medical Education, Inc., for providing the artwork. The authors also thank Mitsuaki Takemi, Tatsuro Fujimaki, Yoshihito Muraoka, and Toko Haga for their technical support, Shoko Tonomoto, Aya Kamiya, and Sayoko Ishii for their general support, and Kurumi Adachi for proofreading support.

