## Supplementary Information for "Closed-loop control of sensorimotor aperiodic neural dynamics can manipulate excitatory-inhibitory balance in humans"

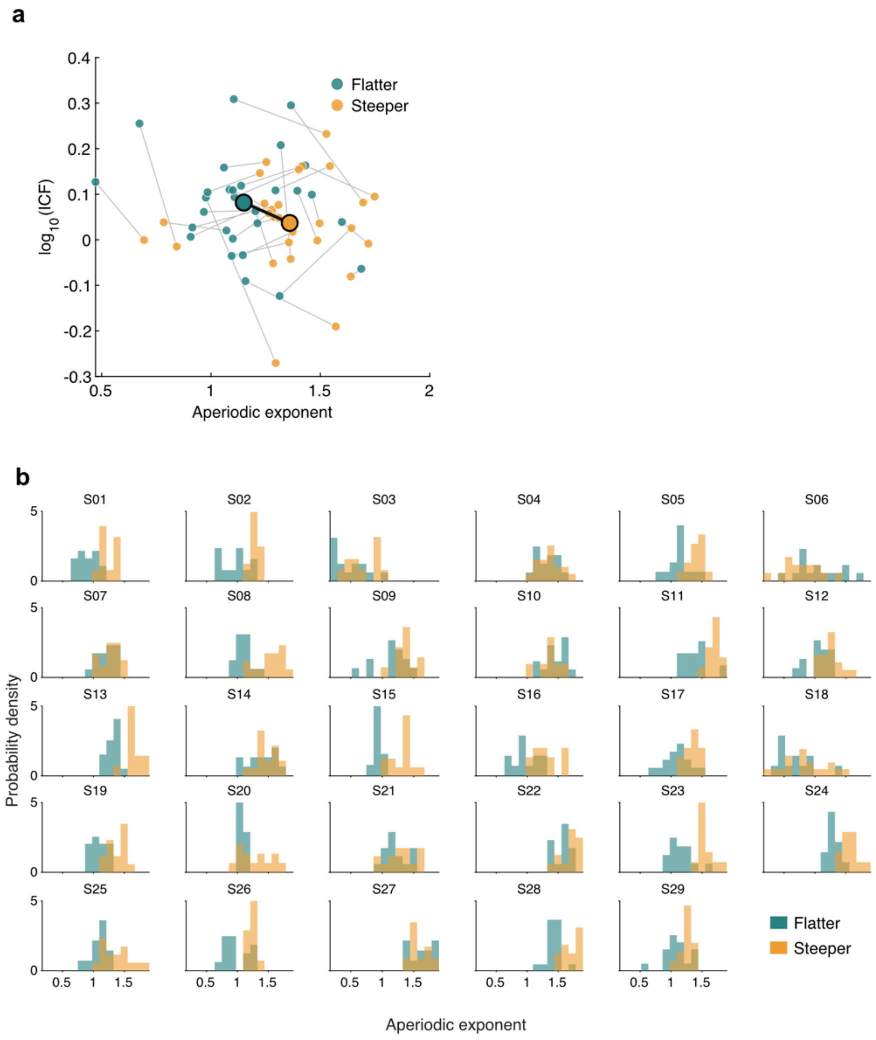


**Supplementary Figure 1. Condition- and trial-level distributions of the aperiodic exponent across neurofeedback states.** (a) Subject-level mean aperiodic exponent and mean log10-transformed ICF responses in the flatter and steeper neurofeedback states. Each pair of points represents the condition means for an individual participant, with paired observations connected by gray lines. Large circles indicate the group means, and the black line indicates the group-level shift from the flatter to the steeper state. (b) Trial-wise distributions of the aperiodic exponent for each participant, restricted to trials in the ICF condition, in the flatter and steeper neurofeedback states. Histograms show the probability density of trial-wise exponent values within each neurofeedback state. Teal and orange indicate the flatter and steeper states, respectively.

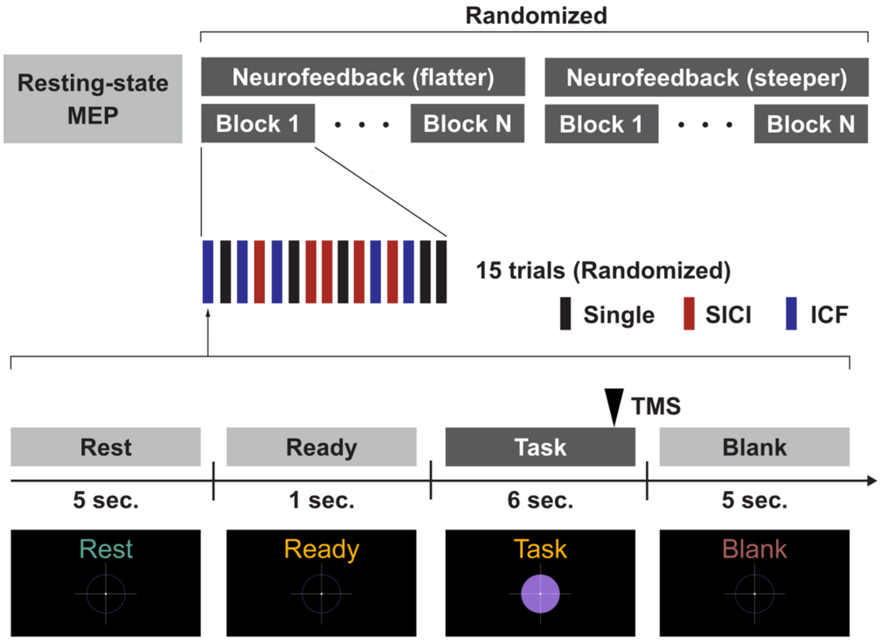


**Supplementary Figure 2. Experimental paradigm.** Following resting-state TMS measurements, participants completed two neurofeedback conditions targeting either a flatter or steeper sensorimotor EEG aperiodic state. The order of the two neurofeedback conditions was randomized across participants. Each neurofeedback condition consisted of successive blocks of 15 trials and continued until 12 successfully triggered trials had been obtained for each of the three TMS conditions (single-pulse TMS, SICI, and ICF), with a maximum of 12 blocks. In the first block, five trials each of single-pulse TMS, SICI, and ICF were presented in randomized order. From the second block onward, the allocation of the three TMS conditions was adaptively randomized according to the accumulated number of successfully triggered trials for each stimulation type. Each trial comprised rest (5 s), ready (1 s), task (6 s), and blank (5 s) phases. During the task phase, participants attempted to modulate the aperiodic exponent in the target direction using real-time visual feedback. TMS triggering began 1 s after task onset and occurred when the feedback signal reached the predefined target threshold (−15% for flatter and +15% for steeper). If the threshold was not reached, TMS was delivered at a randomly selected fallback time during the final 1 s of the task phase.

**Supplementary Movie 1. Validation of threshold-crossing stimulus triggering using a simulated EEG signal.** The movie demonstrates the temporal performance of the closed-loop EEG–TMS system using simulated EEG signals designed to produce either flatter or steeper aperiodic exponents. The simulated signals were fed into the system to evaluate the delay between the aperiodic exponent reaching the predefined modulation threshold and the triggering of TMS.
